# Single-cell Spatial Transcriptomics Reveals Disease-specific Microenvironmental Niches in Neurodegeneration and COVID-19

**DOI:** 10.1101/2025.03.15.643484

**Authors:** Simon Graf, Nicole Ludwig, Matthias Flotho, Annika Engel, Ian F. Diks, Paula Zimmer, Wiebke Jürgens-Wemheuer, Arne Wrede, Daniela Mirzac, Gabriel Gonzalez-Escamilla, Victoria Wagner, Vilas Menon, Sergiu Groppa, Philip L. De Jager, Lai Yiwei, Miguel A. Esteban, Walter J. Schulz-Schaeffer, Andreas Keller

**Affiliations:** Clinical Bioinformatics, Saarland University, 66123, Saarbrücken, Germany; Helmholtz Institute for Pharmaceutical Research Saarland (HIPS)–Helmholtz Centre for Infection Research (HZI), Saarland University Campus, 66123, Saarbrücken, Germany; BGI Research, Shenzhen, China; Department of Human Genetics, Saarland University, 66421 Homburg, Germany; Institute of Neuropathology, Faculty of Medicine, Saarland University, 66421 Homburg, Germany; Department of Neurology, Faculty of Medicine, Saarland University, 66421 Homburg, Germany; Department of Neurology and Neurological Sciences, Stanford University, Stanford, CA, 94305, USA; Center for Translational and Computational Neuroimmunology, Department of Neurology, Columbia University Irving Medical Center, New York, NY, 10032, USA

**Author notes:** Correspondence should be addressed to Andreas Keller and Simon Graf. These authors contributed equally (Walter J. Schulz-Schaeffer, Andreas Keller).

## Abstract

Neurodegenerative diseases and infections can produce lasting effects on brain function, yet the spatial molecular mechanisms underlying these changes remain poorly understood. Here, we present high-resolution spatial transcriptomics of 40 postmortem brain samples from patients with Parkinson’s disease, frontotemporal dementia, dementia with Lewy bodies, and severe COVID-19. Analyzing over 1.5 million spatially resolved cells across dorsolateral prefrontal cortex and anterior cingulate cortex revealed disease-specific transcriptional signatures with pronounced layer-and region-specificity. In Parkinson’s disease, we identified stressed neurons creating distinctive microenvironmental gradients where metabolic and protein degradation pathways are elevated near stress epicenters, while regenerative processes increase with distance. COVID-19 brains displayed extensive peripheral immune cell infiltration, particularly in the subcortical white matter, accompanied by compromised blood-brain barrier and coordinated neuroinflammatory responses from microglia, astrocytes, and endothelial cells. Integration of miRNA sequencing with spatial transcriptomics uncovered layer-specific regulatory patterns, including neuroinflammation-associated miR-155. This atlas provides unprecedented insights into disease pathology and highlights the critical importance of spatial molecular context in understanding brain disorders.

**Key Messages:**

[1] A high-resolution single-cell spatial transcriptomics atlas of the dorsolateral prefrontal cortex and anterior cingulate cortex across neurodegenerative conditions and severe COVID-19
[2] Region-and layer-specific transcriptional dysregulation across disease comparisons reveals disease-specific differential vulnerability
[3] Metabolically stressed cells found selectively in the anterior cingulate cortex of Parkinson’s disease patients but not in dementia with Lewy bodies, along with detailed characterization of their spatial microenvironment
[4] Peripheral immune cell clusters identified in the white matter of the cerebral cortex of COVID-19 patients, with detailed characterization of their spatial microenvironment
[5] Integration of bulk miRNA sequencing reveals cortical layer-specific miRNA regulatory patterns

## INTRODUCTION

Many diseases significantly affect human health, including both communicable and non-communicable conditions. However, these categories are not strictly separate, as infections can trigger long-term health consequences that persist for years or even decades. This phenomenon, often referred to as post-acute infection syndrome (PAIS), is exemplified by the well-established link between viral infections and neurodegenerative diseases. For instance, Herpes simplex virus type 1 (HSV-1) has been implicated in the development of Alzheimer’s disease^1^, with viral DNA detected in amyloid plaques, and positive HSV antibody tests are correlated with worse MMSE (mini mental state examination) outcomes^2^. Similarly, Epstein-Barr virus (EBV) is a risk factor for multiple sclerosis^3,4^. COVID-19, caused by SARS-CoV-2, is associated with long-term complications, collectively referred to as long COVID, which can affect multiple organ systems and persist well beyond the acute infection phase^5–9^. In single-nucleus RNA sequencing (snRNA-seq) of human brain samples from patients who died with severe infections, we identified molecular changes mimicking those seen in neurodegenerative disorders^10^. These examples illustrate how infections can contribute to chronic and systemic health conditions, even long after the initial illness has resolved.

Despite growing evidence linking infections to long-term neurological consequences, the spatial and cellular processes driving these effects are largely unresolved. Understanding how infections, including SARS-CoV-2, contribute to neurodegenerative diseases like Parkinson’s disease (PD), frontotemporal dementia (FTD), and dementia with Lewy bodies (DLB) at a molecular level is crucial for identifying potential therapeutic targets and preventive strategies. These neurodegenerative diseases represent a spectrum of conditions with distinct clinical presentations and neuropathological hallmarks. PD is characterized by the progressive dysfunction and subsequent loss of dopaminergic neurons in the substantia nigra and the presence of synaptic α-synuclein aggregates as well as their aggresome-like concentration in form of Lewy bodies, manifesting clinically as motor symptoms including tremor, rigidity, and bradykinesia, often accompanied by cognitive decline in later stages^11–13^. DLB shares the α-synuclein pathology of PD but presents primarily with cognitive fluctuations, visual hallucinations, and parkinsonism, with synaptic α-synuclein aggregates and Lewy bodies additionally distributed throughout cortical regions^14,15^. FTD encompasses a heterogeneous group of disorders featuring progressive degeneration of frontal and temporal lobes, clinically manifesting as behavioral changes, language deficits, or motor dysfunction, with diverse molecular pathologies including tau, TDP-43, or FUS protein aggregates^16^. The neurological manifestations of COVID-19 have emerged as a significant concern, with patients experiencing various neurological symptoms ranging from nausea and headache to encephalopathy, vascular disorders, and long-term cognitive deficits, though the underlying neuropathological mechanisms remain incompletely understood^5–7,17–20^. Despite their clinical and pathological distinctions, these neurodegenerative conditions share common features of neuroinflammation, cellular stress, and regional vulnerability within the brain^10,21,22^. However, how these processes manifest at the cellular and microenvironmental levels, and how they differ between conditions, remains poorly understood.

Traditional approaches to studying brain pathology have relied on bulk tissue analysis or dissociated single-cell techniques^10,23–26^, which either obscure cellular heterogeneity or lose critical spatial context essential for understanding intercellular interactions and microenvironmental influences. Recent advances in spatial transcriptomics technologies have enabled the simultaneous analysis of gene expression and spatial organization within intact tissue, initially pioneered in model organisms such as mice^27–30^, macaque^31,32^, and axolotl^33^, and now offering unprecedented insights into the complex cellular architecture of the human brain^34–38^. Additionally, microRNAs (miRNAs) have emerged as critical post-transcriptional regulators of gene expression in the central nervous system, with recent large-scale initiatives revealing their importance in neurodegenerative conditions like Parkinson’s disease.^39^ Understanding how these regulatory molecules function within the spatial context of the brain is essential, as their activity may contribute to the layer-specific vulnerabilities observed in these diseases. Together, the ability to precisely map both transcriptional changes and miRNA regulation to specific brain regions, cortical layers, and cellular neighborhoods could provide crucial insights into disease mechanisms and potentially identify novel therapeutic targets.

Here, we present a comprehensive analysis and comparison of high-resolution single-cell spatial transcriptomics and paired miRNA sequencing data from two cortical regions, the dorsolateral prefrontal cortex (DLPFC) and anterior cingulate cortex (ACC), across patients with PD, FTD, DLB, and severe COVID-19. This approach allowed us to characterize disease-specific transcriptional signatures within distinct cortical layers and white matter, identify stressed cell populations and their surrounding microenvironments in PD, characterize mechanisms of peripheral immune cell infiltration in COVID-19 brains, and identify layer-specific miRNA regulation profiles.

## RESULTS

### Single-cell spatial transcriptomics of the human neocortex indicates consistent cytoarchitecture across conditions

We performed high-resolution spatial transcriptomics on brain sections from patients with neurodegenerative conditions and severe COVID-19 using the Stereo-seq platform^27^. This platform enables high spatial resolution (500×500 nm center-to-center distance) at both, the spot-level (200×200 Stereo-seq spots, bin200) and for single-cell-level analyses. Thereby, it captures both anatomical and cellular characteristics (see Methods). Our cohort includes 40 postmortem human brain samples, comprising patients with Parkinson’s disease (PD; n=5), frontotemporal dementia (FTD; n=4), dementia with Lewy bodies (DLB; n=6), and severe COVID-19 (n=5). For each patient, we selected and compared two brain regions: the dorsolateral prefrontal cortex (DLPFC; Brodmann area 9,46) and the anterior cingulate cortex (ACC; Brodmann area 33), yielding a total of 40 Stereo-seq samples (**Fig. 1a**). Following quality control based on tissue integrity and expression of established cortical layer markers (MBP, SNAP25, PCP4), 35 high-quality samples were retained for subsequent analysis. Comprehensive patient characteristics, including age, sex, post-mortem interval and cause of death, are provided in **Supplementary Table 1**. The final dataset comprised more than 150,000 spots and 1.5 million spatially resolved single cells (**Fig. 1b**), with no significant differences in spot or cell numbers observed across diseases or brain regions.

**Figure 1:**
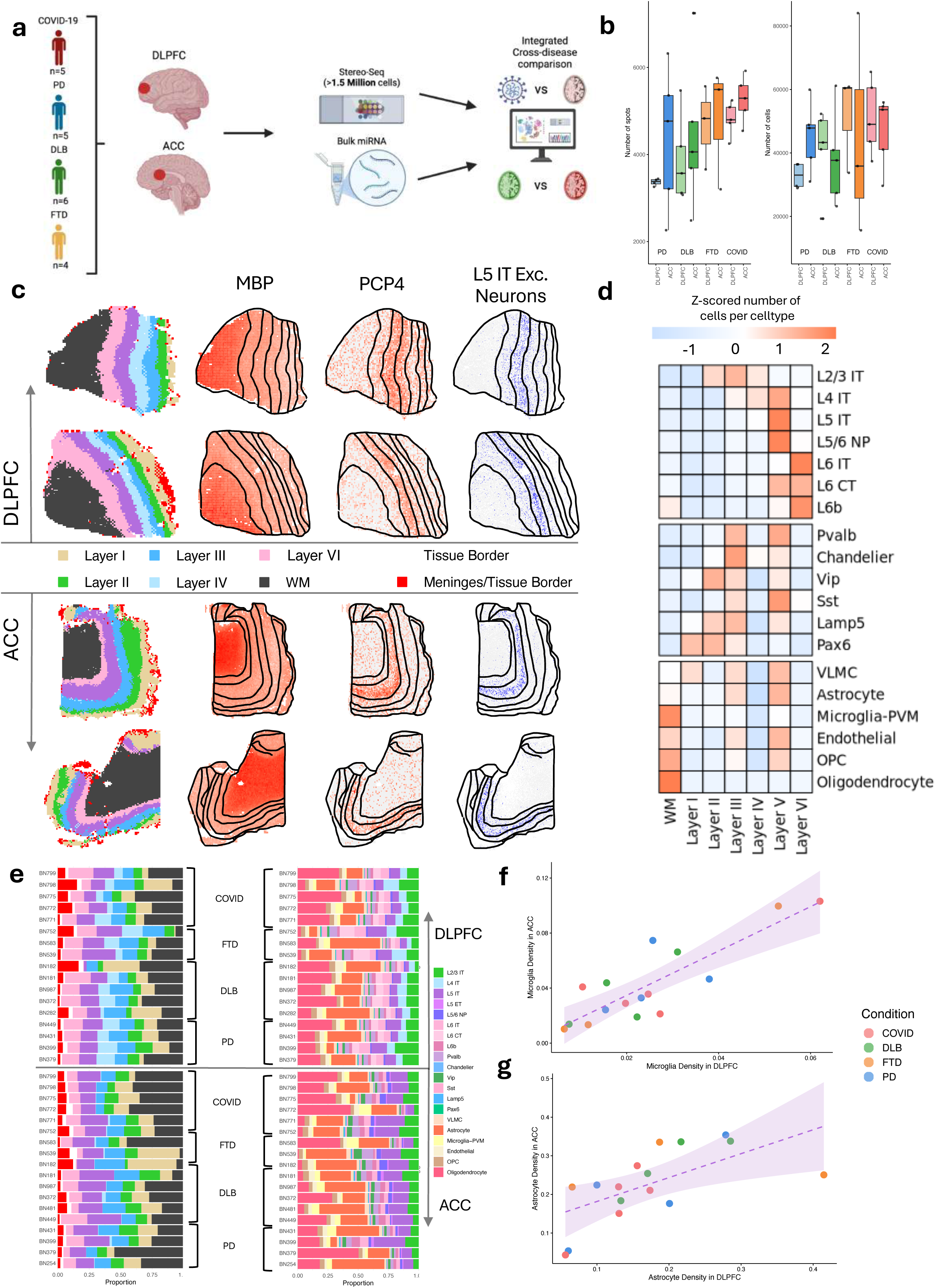
Spatial transcriptomics of the human neocortex **a,** Schematic illustration of the experimental workflow and study design. **b,** Boxplots depicting the distribution of spot and cell counts across experimental groups. **c,** Visualization of spatial domain clustering and annotation, alongside marker genes MBP and PCP4 within tissue, exemplifying cytoarchitecturally consistent cell type annotation with L5 IT neurons. Manual contours delineate distinct spatial domains. **d,** Heatmap displaying z-scored cell type distribution across spatial domains. **e,** Bar chart illustrating the composition of spatial domains and cell types for each sample. **f,** Scatter plot demonstrating the correlation of microglia density between ACC and DLPFC regions. **e,** Scatter plot demonstrating the correlation of astrocyte density between ACC and DLPFC regions.

We employed BayesSpace^40^ for spatial domain segmentation on spot-based data to classify each spot into white matter or cortical layers 1-6 based on gene expression profiles and spatial positioning (see **Methods**). Expression patterns of established anatomical marker genes aligned with our segmented domains; for instance, MBP, a major oligodendrocyte marker, showed highest expression in white matter areas, while PCP4 was predominantly expressed in cortical layer V, consistent with previous reports^34,41^ (**Fig. 1c**). We then applied DestVI^42^ to transfer cell type annotations from the SeattleAD atlas^43^ onto our spatially resolved single cells (see **Methods**), identifying all major glial cell populations as well as excitatory and inhibitory neuron subtypes (**Fig. 1c,d**). All identified cell types expressed canonical marker genes previously reported in the literature, including MBP, GFAP, CD74 and SNAP25 (**Supplementary Fig. 1a-c**). We subsequently mapped the domain annotations from spots to corresponding single cells to obtain anatomical labels for each cell, enabling quantification of anatomical distribution patterns for each cell type (**Fig. 1c,d**). The observed distributions aligned with established knowledge of brain cytoarchitecture; for instance, oligodendrocytes were predominantly localized in white matter, while L5 neuron subtypes were mainly found in cortical layer 5.

The anatomical domain and cell type compositions were comparable across all samples, with no significant differences observed between disease groups (**Fig. 1e,f**). Notably, we identified virtually no spots in the ACC classified as layer IV and no cells classified as L4 IT neurons, consistent with the established cytoarchitectural feature that the ACC lacks a prominent layer IV^44^. Correlating cell type densities of the ACC and DLPFC of the same patient shows strong correlation in microglia and astrocytes (microglia: p<10^-5^, R=0.854; astrocytes: p<0.01, R=0.628). pointing to potentially patient-specific glial responses with no clear patterns across disease groups (**Fig. 1g,h**). Interestingly, we observed that microglia density was approximately twofold higher in the ACC compared to the DLPFC across all samples, suggesting regionally elevated neuroimmune activity independent of disease pathology. In summary, we established a high-resolution spatial transcriptomic atlas spanning two cortical regions in neurodegenerative and COVID-19 brain samples, incorporating both spot-based anatomical and cell-based perspectives, while observing consistency in fundamental anatomical and cellular composition across diseases. To leverage this atlas and systematically investigate transcriptional differences across patients, spatial domains, and disease groups, we now perform a thorough cross-disease analysis.

### Global spatial domain comparison indicates layer-and region-specific differences

We implemented a pseudobulk procedure that generated aggregated expression profiles for each unique combination of patient, spatial domain, disease type, and brain region (see **Methods**). Following standard preprocessing and normalization, we performed dimensionality reduction using UMAP to project these composite transcriptional profiles into two-dimensional space (**Fig. 2a**). The resulting embedding revealed clear transcriptional segregation between the ACC and DLPFC regions. Within these regional clusters, distinct groupings corresponding to different cortical domains were evident. White matter and Layer 1 exhibited the most pronounced separation from other domains, likely reflecting their distinctive cellular compositions compared to cortical layers 2-6. Within each domain cluster, we further observed subclustering by disease group, highlighting disease-specific transcriptional signatures that persisted across spatial domains.

**Figure 2:**
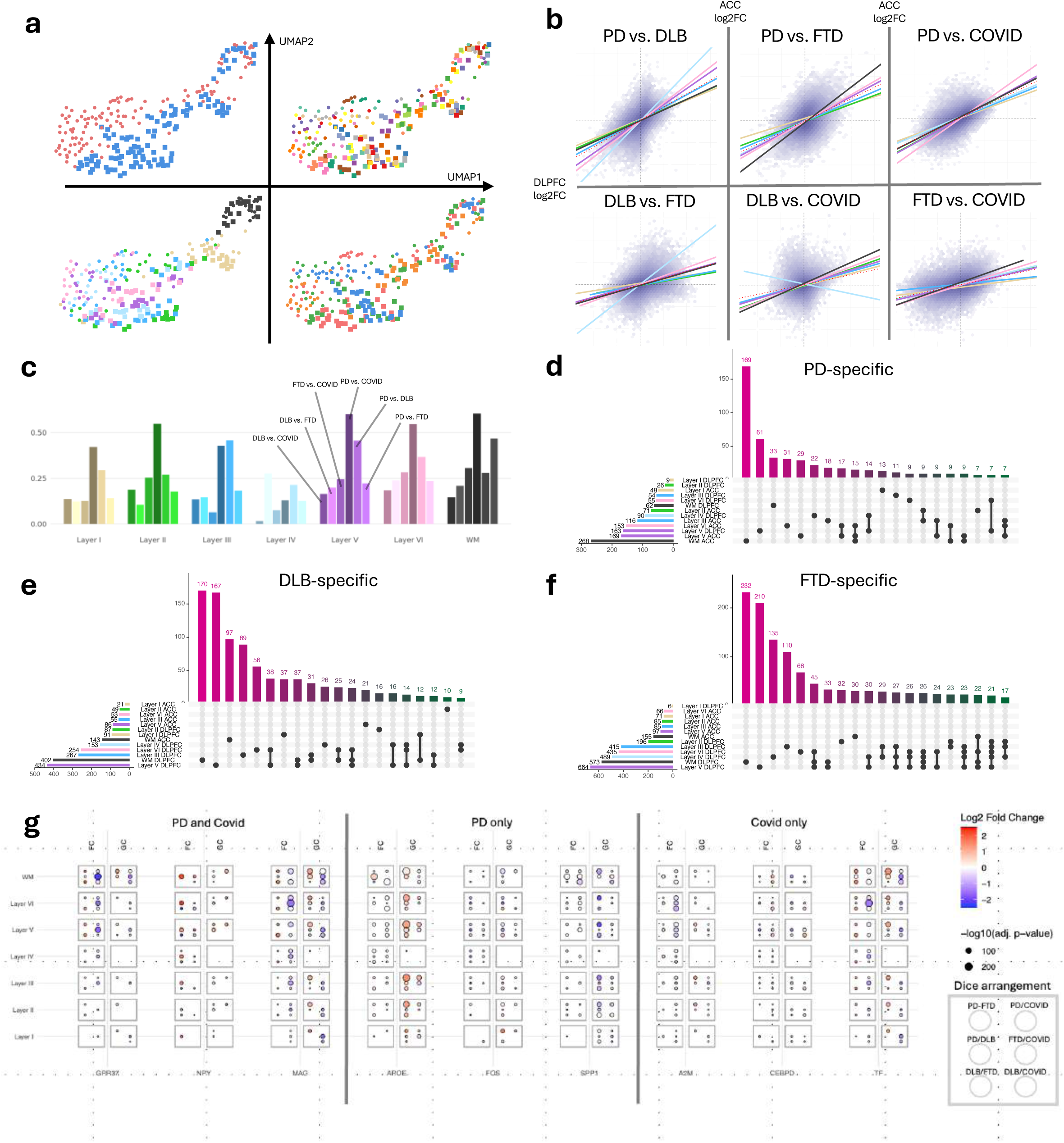
Global domain comparison across diseases **a,** 2D UMAP embedding generated from pseudobulk samples across region, domain, and patient. Embeddings are color-coded by region, patient, domain, and condition. Circular markers represent DLPFC samples while square markers indicate ACC samples. **b,** Density plots illustrating log2 fold change correlation between ACC and DLPFC for all pairwise disease comparisons. Different colored lines represent distinct spatial domains. **c,** Bar chart displaying R² values for each comparison across all spatial domains. Different shades within each domain represent the six distinct disease comparisons. **d,** Upset plot visualizing the intersection of PD-specific differentially expressed gene (DEG) sets across spatial domains and cortical regions. **e,** Upset plot visualizing the intersection of DLB-specific differentially expressed gene (DEG) sets across spatial domains and cortical regions. **f,** Upset plot visualizing the intersection of FTD-specific differentially expressed gene (DEG) sets across spatial domains and cortical regions. **g,** Dice plot illustrating the dysregulation patterns of specific gene sets: those dysregulated in both PD and COVID-19, those unique to PD, and those unique to COVID-19, across six comparisons and both brain regions.

To get an in-depth understanding of disease-specific effects, we performed pairwise differential expression analysis between all disease groups, stratified by spatial domain (see **Methods**) (**Supplementary Table 2**). To evaluate transcriptional consistency between the ACC and DLPFC, we calculated correlations between fold changes for shared genes and domain combinations across all six disease-pair comparisons. Across all diseases, we observed positive and significant correlations between the two brain regions, indicating substantial transcriptional similarities in disease-associated gene expression patterns (**Fig. 2b**). The strength of these region-to-region correlations varied significantly across cortical layers: layer 5 and the white matter present the strongest correlations, while cortical layer 1 and layer 4 display weaker correlation (**Fig. 2b,c**). The particularly weak correlation in layer 4 aligns with known cytoarchitectural differences between ACC and DLPFC, with ACC featuring a less prominent layer 4^44^. Among disease comparisons, PD versus DLB and PD versus COVID-19 consistently showed the highest correlations across all spatial domains (**Fig. 2b,c)**, suggesting greater regional transcriptional similarity in those comparisons.

To identify transcriptional signatures unique to each condition, we defined disease-specific differentially expressed genes (DEGs) as those significantly up-or down-regulated in one disease compared to at least two other diseases. We then examined the overlap of these disease-specific DEGs across brain regions and cortical layers. The majority of DEGs exhibited marked layer-and region-specificity, with minimal overlap between different layers of the same region and virtually no shared DEGs between the ACC and DLPFC (**Fig. 2d-f, Supplementary Fig. 2a**). Intriguingly, despite the substantial positive correlation of fold changes between regions, we observed few common DEGs. This pattern suggests that while most genes may be regulated in the same direction across regions, they frequently fail to meet statistical significance thresholds in the other region, highlighting the importance of regional context in disease-specific transcriptional dysregulation. Of note, each disease exhibited distinctive regional and domain preferences in transcriptional dysregulation (**Fig. 2d-f, Supplementary Fig. 2a**). FTD and DLB showed predominant dysregulation in the DLPFC, while PD displayed a more balanced pattern with stronger dysregulation in the ACC. COVID-19 generally showed greater dysregulation in the DLPFC; however, the ACC white matter exhibited exceptionally strong transcriptional alterations compared to other ACC domains. As infectious diseases like COVID-19 are increasingly recognized to mimic features of neurodegenerative diseases, we next aimed to compare transcriptional dysregulation between COVID-19 and PD. For both diseases, we performed differential expression analysis against FTD and DLB, yielding both overlapping and disease-specific DEGs. We employed a dice plot^45^ to demonstrate that both overlapping and disease-specific DEGs exhibit clear layer-and region-specific dysregulation patterns (**Fig. 2g**)

In summary, our analysis revealed comprehensive transcriptional dysregulation exhibiting distinct layer-dependent and region-specific patterns, highlighting unique spatial vulnerability signatures characteristic of the disease pathology.

### Characterizing the microenvironment of stressed cells in PD

Despite their shared α-synuclein aggregate pathology, Parkinson’s disease (PD) and dementia with Lewy bodies (DLB) present distinct clinical and neuropathological features^46^. Given the strong transcriptional dysregulation between both diseases observed at the anatomical level in our previous analyses, we performed in-depth molecular characterization at single-cell resolution to elucidate the underlying differences between these α-synuclein aggregation diseases in more detail. Differential gene expression analysis of neuronal populations revealed a pronounced upregulation of stress response pathways in PD compared to DLB in the ACC, but not in the DLPFC (**Fig. 3a**, Supplementary Figure 3a, Supplemental Table 3).

**Figure 3:**
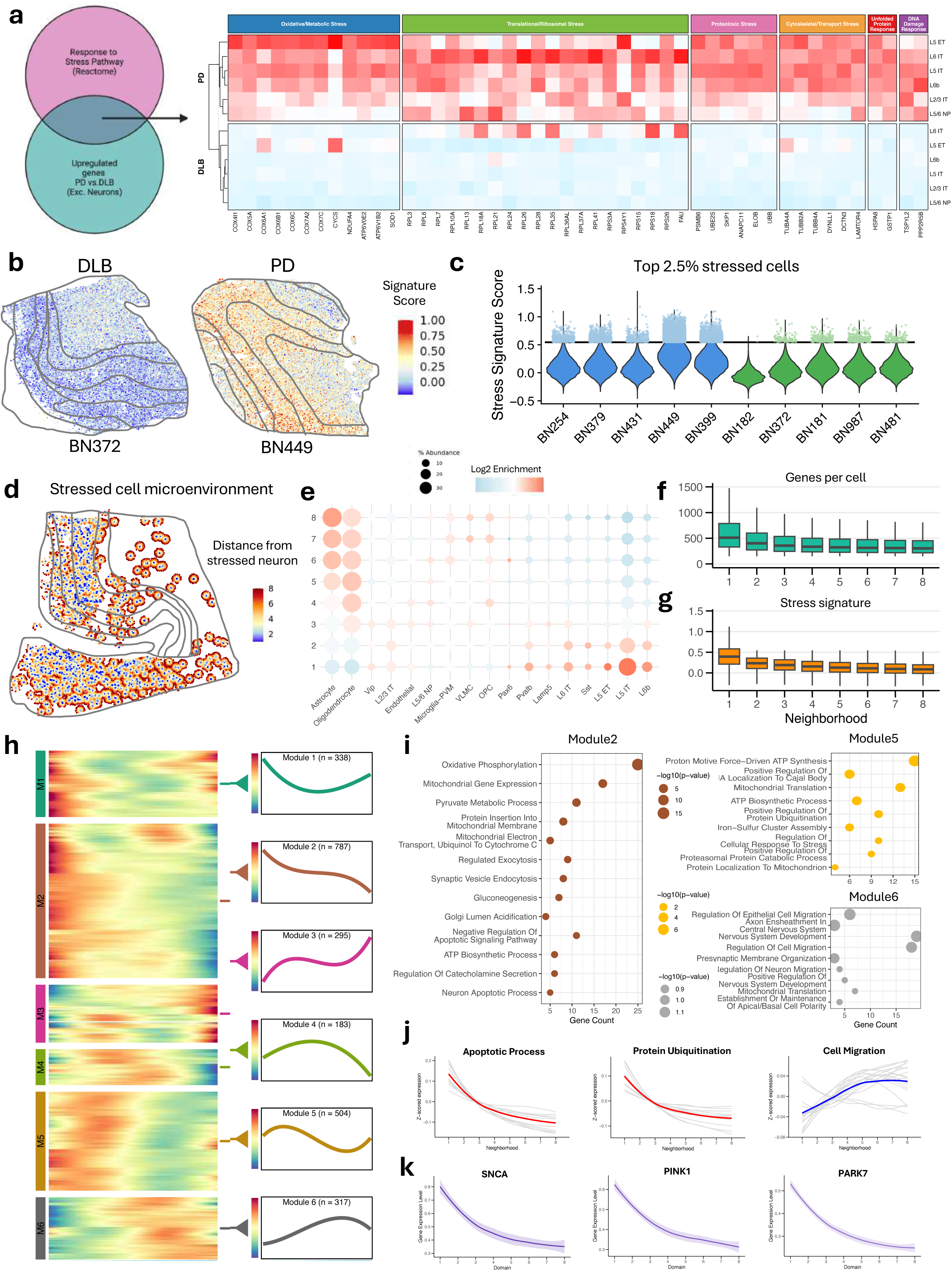
Characterizing the microenvironment of PD stressed cells **a,** Compositional overview of the stress signature and heatmap displaying z-scored average expression profiles in ACC excitatory neurons, comparing PD and DLB patients. **b,** Spatial visualization contrasting stress signature scores between representative PD and DLB samples. Manual contours delineate distinct spatial domains based on domain segmentation analysis. **c,** Violin plot comparing stress signature scores across all patients, with PD (blue) and DLB (green) differentiation. The horizontal threshold identifies the top 2.5% of cells with highest signature scores, classified as stressed cells. **d,** Spatial visualization highlighting cells in neighborhoods 1-8, with all other cells omitted for clarity. **e,** Dotplot representing cell type distribution across neighborhoods 1-8. Dot size indicates relative abundance within each neighborhood, while color intensity denotes log2 enrichment scores for each cell type. **f,** Boxplot showing stress signature distribution across neighborhoods 1-8. **g,** Boxplot showing stress signature distribution across neighborhoods 1-8. **h,** Heatmap and expression trajectory illustrating the expression patterns of modules 1-6 across neighborhoods 1-8. **i,** Dotplot summarizing pathway enrichment analysis results for genes from modules 2, 5, and 6. **j,** Line plot tracking the trajectory of GO pathways Negative regulation of apoptotic process, Protein Ubiquitination, and Cell Migration across neighborhoods 1-8. Gray lines represent individual gene trajectories, while colored lines show mean expression across all pathway genes per neighborhood. Only genes within the enriched gene sets are displayed. **k,** Line plot depicting normalized expression patterns of SCNA, PINK1, and PARK7 genes across neighborhoods 1-8.

To systematically identify cells exhibiting stress phenotypes, we generated a stress response gene signature by intersecting upregulated genes in PD excitatory neurons with the Reactome “Cellular response to stress” pathway^47^. This signature contains 42 genes, most of them related to metabolism (ATP6V0E2, NDUFA4), oxidative phosphorylation (COX4I1, COX5A), RNA translation (RPL3, RPL6) and proteasomal components (PSMB6, UBB), recapitulating established molecular hallmarks of PD pathophysiology^48^. Quantification of this signature across individual cells demonstrated significantly elevated stress response in PD relative to DLB sample (**Fig. 3b,c**). We classified the top 2.5% of cells with the highest signature scores as “stressed” cells for subsequent analyses. While substantial inter-individual variability in stressed cell abundance was observed within the PD cohort, the number of stressed cells per donor was significantly higher in PD compared to DLB patients (Figure 2d). Two patients (BN399, BN449) showed especially high numbers of stressed cells.

To understand both the localized reactions to stressed cells and the influence these cells exert on their surroundings, we next sought to characterize their spatial microenvironment. We implemented a distance-based classification system wherein each cell was assigned a neighborhood label based on its proximity to the nearest stressed cell (**Figure 3d**). Specifically, stressed cells and their immediate surrounding cells were designated as neighborhood 1, with the neighborhood designation incrementing at 50μm intervals from stressed cells (see **Methods**). This spatial stratification enabled systematic analysis of transcriptional gradients and cellular interactions as a function of distance from stressed cell populations. We quantified cell type composition across neighborhoods to identify populations enriched or depleted in proximity to stressed cells (**Fig. 3e**). As anticipated, subgranular excitatory neuron populations (L5 ET, L5 IT, L6 IT, L6b) were significantly enriched within and immediately adjacent to stressed cells. In contrast, glial lineages, particularly astrocytes and oligodendrocytes, exhibited relative depletion in the immediate microenvironment of stressed cells, with their proportional representation increasing as a function of distance. Notably, despite this relative depletion, astrocytes and oligodendrocytes remained substantial constituents of the stressed cell microenvironment due to their overall abundance in the tissue. As expected, the stress signature scores were highest in neighborhood 1 and decreased with increasing distance from stressed cells (**Fig. 3f**). Interestingly, we discovered that the number of genes expressed per cell was also highest in stressed cells and gradually diminished with distance from these cells (**Fig. 3g)**. This spatial gradient in transcriptional output suggests a compensatory increase in translational machinery at the epicenter of cellular stress, which gradually tapers off in the surrounding tissue microenvironment. To comprehensively characterize the transcriptional landscape across neighborhoods, we conducted a holistic gene expression gradient analysis. We first identified genes with significant expression differences across neighborhoods and then clustered them into modules based on their transcriptional trajectories (**Fig. 3h**). Module 2 and Module 5 comprised genes with elevated expression in stressed cells and their immediate microenvironment that progressively decreased with distance. Pathway enrichment analysis of these modules (excluding genes contained in the stress signature) revealed significant overrepresentation of metabolic pathways, as well as apoptotic and ubiquitination processes (**Fig. 3i,j**). Conversely, Module 6 consisted of genes with reduced expression in stressed cells that gradually increased with distance, showing enrichment primarily for regenerative pathways including cell migration and nervous system development.

These spatial transcriptional gradients suggest a stress-response continuum in the PD cortex. The enrichment of metabolic, apoptotic, and protein degradation pathways proximal to stressed neurons likely reflects cellular attempts to maintain homeostasis under stress conditions. Meanwhile, the inverse gradient of regenerative pathways increasing with distance from stressed cells may represent compensatory neuroprotective mechanisms activated in less affected cells. Intriguingly, we see that prominent PD risk genes (SNCA, PINK1 and PARK7) show a clear expression gradient being approximately twofold higher expressed in stressed cells and their immediate neighborhood than in neighborhood 8 (**Fig. 3k**).

In summary, we identified a specific population of stressed cells that are significantly more abundant in individuals with PD than with DLB, which create a distinctive microenvironmental gradient where stress-response pathways diminish and regenerative processes increase with distance from these cellular stress epicenters. To better understand the parallels and distinctions between neurodegenerative diseases like PD or DLB and COVID-19, we will now compare the molecular and spatial signatures of neuroinflammation across these conditions, with particular emphasis on the characteristic non-neural responses observed in the subcortical white matter.

### Immune infiltration into the subcortical WM in COVID-19

Although SARS-CoV-2 is not traditionally classified as a neurological disease, recent molecular studies show that it infiltrates the brain, disrupts neural networks, and triggers chronic inflammation—processes that could ultimately contribute to neurodegenerative diseases such as Parkinson’s and dementia ^9,10,18–20,49^. We therefore sought to spatially characterize cortical responses specific to severe COVID-19 that are either absent or markedly attenuated in neurodegenerative diseases. Our previous analyses identified the cortical white matter in both the ACC and DLPFC as particularly vulnerable regions, prompting a more detailed characterization of these domains at single-cell resolution.

Overall, our findings reveal that COVID-19 triggers a distinct pattern of peripheral immune cell infiltration, seen in the white matter of the anterior cingulate cortex (ACC). This infiltration appears to result from a compromised blood-brain barrier (BBB), characterized by endothelial inflammation and activation of both microglia and astrocytes (**Fig. 4a**).

**Figure 4:**
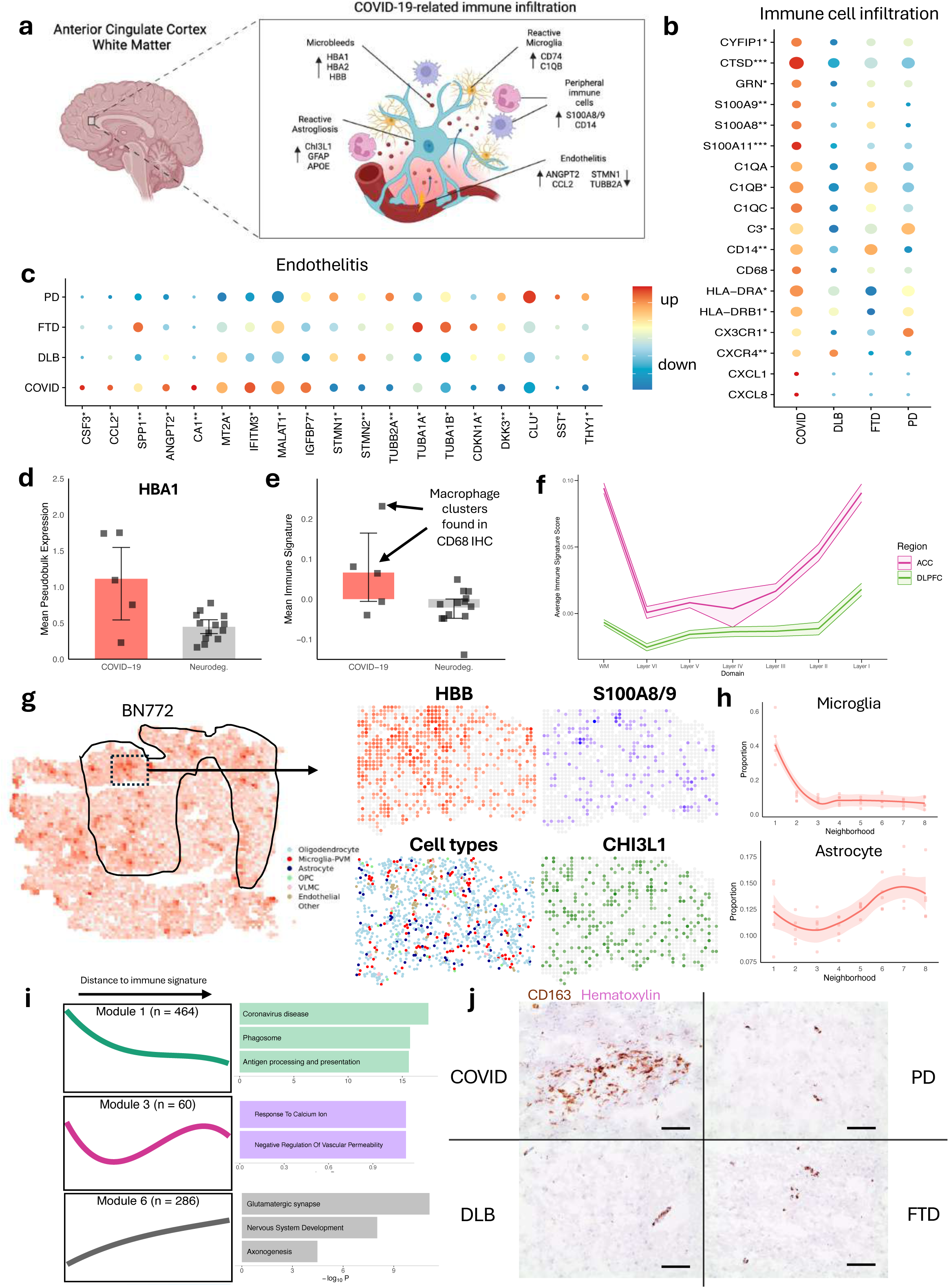
Peripheral Immune Infiltration in the COVID-19 WM **a,** Schematic illustration depicting the proposed COVID-19-related immune infiltration into the subcortical white matter. **b,** Differential gene expression in microglia across all four diseases, specifically within the ACC white matter. The number of asterisks indicates the number of disease comparisons in which genes show significant dysregulation relative to COVID-19. **c,** Differential gene expression in endothelial cells across all four diseases, specifically within the ACC white matter. The number of asterisks indicates the number of disease comparisons in which genes show significant dysregulation relative to COVID-19. **d,** Boxplot comparing HBA1 pseudobulk expression per sample in the ACC white matter between COVID-19 and the pooled cohort of all three neurodegenerative conditions. **e,** Boxplot comparing the mean immune signature in the ACC white matter per patient between COVID-19 and the pooled cohort of all three neurodegenerative conditions. **f,** Distribution of immune signature across all spatial domains, comparing the ACC and the DLPFC. **g,** Spatial visualization showing HBB expression in tissue (bin50). Inset displays spatial expression patterns of HBB, combined S100A8/9 (summed per spot), and CHI3L1, alongside the spatial distribution of cell types. **h,** Cell type proportions of microglia and astrocytes across neighborhoods 1-8 within immune hotspots. **i,** Expression trajectories of modules 1, 3, and 6 across neighborhoods, with accompanying bar charts showing enriched pathways within each corresponding module. **j,** CD163 immunohistochemistry in the ACC from one COVID-19 sample and one representative sample from each of the three neurodegenerative diseases. Scale bars: 20 μm.

We first performed differential expression analysis in three major cell populations (microglia, endothelial cells, and astrocytes) in the cortical white matter to understand their transcriptional responses to COVID-19 infection compared to neurodegenerative conditions (Supplemental Table 4). In the ACC, we identified a substantial set of inflammatory markers upregulated in COVID-19 relative to at least one neurodegenerative disease (**Fig. 4b; Supplementary Table 3**). These included peripheral macrophage/monocyte markers (S100A8, S100A9, CD14)^50^, complement pathway components (C1QB, C3), type II MHC molecules (HLA-DRA, HLA-DRB1), and chemokine receptors (CX3CR1, CXCR4). The upregulation of these genes indicates the presence of peripheral immune cells distinct from resident microglia, though the definitive separation of these populations is limited by the lower capture efficiency of current spatial transcriptomics methods compared to contemporary single-cell approaches. Nevertheless, these findings are strongly corroborated by previous reports in the literature^51^. In endothelial cells, we observed upregulation of genes related to pro-inflammatory responses (CSF3, CCL2, SPP1, ANGPT2)^52–56^ and immune-response modulation (MT2A, IFITM3, MALAT1)^57–60^, alongside downregulation of genes associated with cytoskeletal stability (STMN1, STMN2, TUBB2A, TUBA1A, TUBA1B). (**Fig. 4c**). These transcriptional changes suggest compromised endothelial integrity and blood-brain barrier dysfunction, further supporting the hypothesis of peripheral immune cell infiltration into the brain parenchyma. White matter astrocytes exhibited a distinct transcriptional profile characterized by increased GFAP expression, indicative of reactive astrogliosis^61^ (**Supplementary Fig. 4a**). We also observed upregulation of mitochondrial genes (MT-ND2, MT-ND3, MT-ND5), suggesting elevated metabolic stress, neuroinflammatory markers (CHI3L1, CHI3L2)^62^, and genes associated with vascular permeability (ANGPTL4, AQP1). Additionally, we see increased APOE expression in astrocytes, which has been previously linked to blood-brain barrier impairment^63^, further supports the compromised neurovascular unit integrity in COVID-19. We validated the upregulation of a subset of these genes using an independent bulk RNA-seq dataset from human cortex samples^64^. Notably, S100A8 and S100A9 exhibited strong upregulation in COVID-19 brain tissue, confirming our spatial transcriptomics findings (**Supplementary Fig. 4b**).

Beyond these cell-type specific alterations, we observed a striking upregulation of hemoglobin-related genes (HBA1, HBA2, and HBB) in COVID-19 white matter. This pattern was evident across multiple cell types and in the spot-level analysis. Pseudobulk analysis at the donor level confirmed significant or near-significant upregulation of these genes (**Fig. 4d, Supplementary Fig. 4c**). While these findings further support our hypothesis of blood-brain barrier compromise in COVID-19, we approach this interpretation with caution. Correlation analysis between hemoglobin gene expression and post-mortem interval revealed a small, non-significant correlation, however we cannot completely exclude the possibility that these apparent microbleeds might be related to post-mortem artifacts rather than disease pathology.

To quantitatively assess immune cell infiltration, we developed a signature based on the upregulated inflammatory markers identified in **Fig. 4b**. We calculated signature scores for all microglia in white matter regions across all samples and aggregated these scores at the donor level. This analysis revealed robust upregulation of the immune signature in COVID-19 patients compared to those with neurodegenerative diseases (**Fig. 4e**). However, we observed considerable inter-individual variability. Three COVID-19 patients exhibited strongly elevated immune signatures, while two patients showed levels comparable to those in neurodegenerative conditions. Notably, the key distinguishing feature between high-signature and low-signature COVID-19 patients was the presence of distinct macrophage clusters in white matter regions, as confirmed by histopathological examination. This correlation between our transcriptional signature and histological findings further validates the association between our immune signature and peripheral immune cell infiltration in COVID-19 white matter.

To further validate our immune signature, we applied it to a previously published single-cell RNA-seq dataset comparing COVID-19 and control samples from both choroid plexus and medial frontal cortex^10^. Thereby, we observed significant upregulation of our immune signature in macrophages within the choroid plexus of COVID-19 patients (p<0.0006, Wilcoxon Mann-Whitney test), but not in the medial frontal cortex (p<0.518) (**Supplementary Fig. 5a**). Analysis of patients common to both datasets revealed consistent patterns of signature expression; patient BN772 exhibited strong upregulation of the signature, while patients BN771 and BN775 showed lower overall signature scores (**Supplementary Fig. 5b**), mirroring the heterogeneity observed in our spatial transcriptomics analysis.

Spatial analysis of immune signature distribution across cortical layers revealed that the immune response was most pronounced in white matter regions (**Fig. 4f**). Comparative analysis between brain regions demonstrated that the ACC exhibited substantially stronger immune activation than the DLPFC. Based on this regional vulnerability observed in our dataset, we focused the subsequent analyses on the ACC white matter. This regional specificity aligns with our single-cell RNA-seq validation findings, where the medial frontal cortex similarly showed no significant increase in immune signature expression.

Within the tissue, we observed that S100A8/9, HBB, and CHI3L1 were expressed in discrete spatial clusters without necessarily colocalizing (**Fig. 4g**). Employing an approach similar to our previous stress response analysis, we calculated the immune signature for each cell and defined the top 5% highest-scoring cells as “immune hotspots”. We then classified cells based on their distance to the nearest immune hotspot into neighborhoods (at 50μm intervals) and characterized the spatial microenvironment through the first eight neighborhood strata (**Supplementary Fig. 6a**). Notably, we observed distinct spatial clustering of immune hotspots, particularly in severely affected individuals with histologically confirmed macrophage clusters in white matter (BN772 and BN799). Analysis of cellular composition across neighborhoods revealed that microglia were enriched in immune hotspots and their immediate vicinity, with their proportion gradually decreasing with increasing distance from these hotspots. Conversely, astrocytes showed relative depletion near immune hotspots, with their representation increasing with distance (**Fig. 4h**). Interestingly, endothelial cells, like microglia, were enriched near and inside immune hotspots, further supporting the peripheral origin of these immune clusters (**Supplementary Fig. 6b**).

To understand the molecular mechanisms underlying these hotspots and their spatial microenvironment, we analyzed gene expression gradients by clustering genes into modules based on their expression trajectories across neighborhoods (**Supplementary Fig. 6c**). Module 1, comprising genes with elevated expression in immune hotspots that gradually diminished with increasing distance, showed strong enrichment for inflammatory pathways related to viral response (Phagosome, Antigen Processing and Presentation) (**Fig. 4i**). In contrast, Module 6, characterized by reduced expression in immune hotspots with progressive increase at greater distances, was enriched for protective and regenerative pathways (Glutamatergic Synapse, Nervous System Development, Axonogenesis). This spatial polarization of inflammatory versus neuroprotective programs suggests that immune infiltration in COVID-19 creates distinct microenvironmental zones where inflammation suppresses regenerative processes in the immediate vicinity of immune hotspots.

Finally, CD163 immunohistochemistry confirmed the presence of peripheral immune cell clusters in the ACC white matter of COVID-19 patients, while no comparable clusters were detected in any of the three neurodegenerative disease groups (**Fig. 4j**). This histological validation substantiates our transcriptional findings and further supports our hypothesis of peripheral immune cell infiltration as a distinctive feature of COVID-19 brain pathology against neurodegeneration.

In summary, we provide compelling evidence for peripheral immune infiltration into the brain specific to severe COVID-19, characterizing in detail the spatial organization and molecular features of the affected microenvironment. The major immune hotspots are pronounced in the white matter of the cortex and accompanied by significant activation of astrocytes and endothelial cells, creating distinct zones where inflammatory processes suppress regenerative pathways.

### miR-155 regulates genes in a layer-specific manner

MicroRNAs (miRNAs) function as critical post-transcriptional regulators of gene expression in the central nervous system. Sequencing of genes^65^ and miRNAs^39^ of 5,450 samples from 1,614 patients included in the Parkinson’s Progression Marker Initiative (PPMI) revealed a key role of selected small non-codingRNAs. Given the established transcriptional heterogeneity across cortical layers, we investigated whether miRNA-mediated regulation might also exhibit layer-specific patterns. To address this question, we performed comprehensive bulk miRNA sequencing across all 40 brain samples.

To investigate layer-specific miRNA regulation, we integrated the miRNA sequencing data with our spatially-resolved transcriptomics dataset. We first filtered the dataset to include only those exceeding predefined quality control thresholds, being significantly expressed above the background (c.f. Methods). We then linked these stable expressed miRNAs to their experimentally validated gene targets using the miRTarBase repository^66^. For the spatial transcriptomics data, we aggregated the single cell data to the pseudo-bulk level for each patient and spatial domain to facilitate a direct comparison with the bulk miRNA sequencing results. Specifically, we computed Spearman correlation coefficients for all miRNA-mRNA target interactions (MTI) across samples. After filtering this approach yielded a comprehensive correlation matrix (272 MTIs x 7 domains) revealing miRNA-mRNA regulatory relationships across all seven spatial domains (**Fig. 5a**).

**Figure 5:**
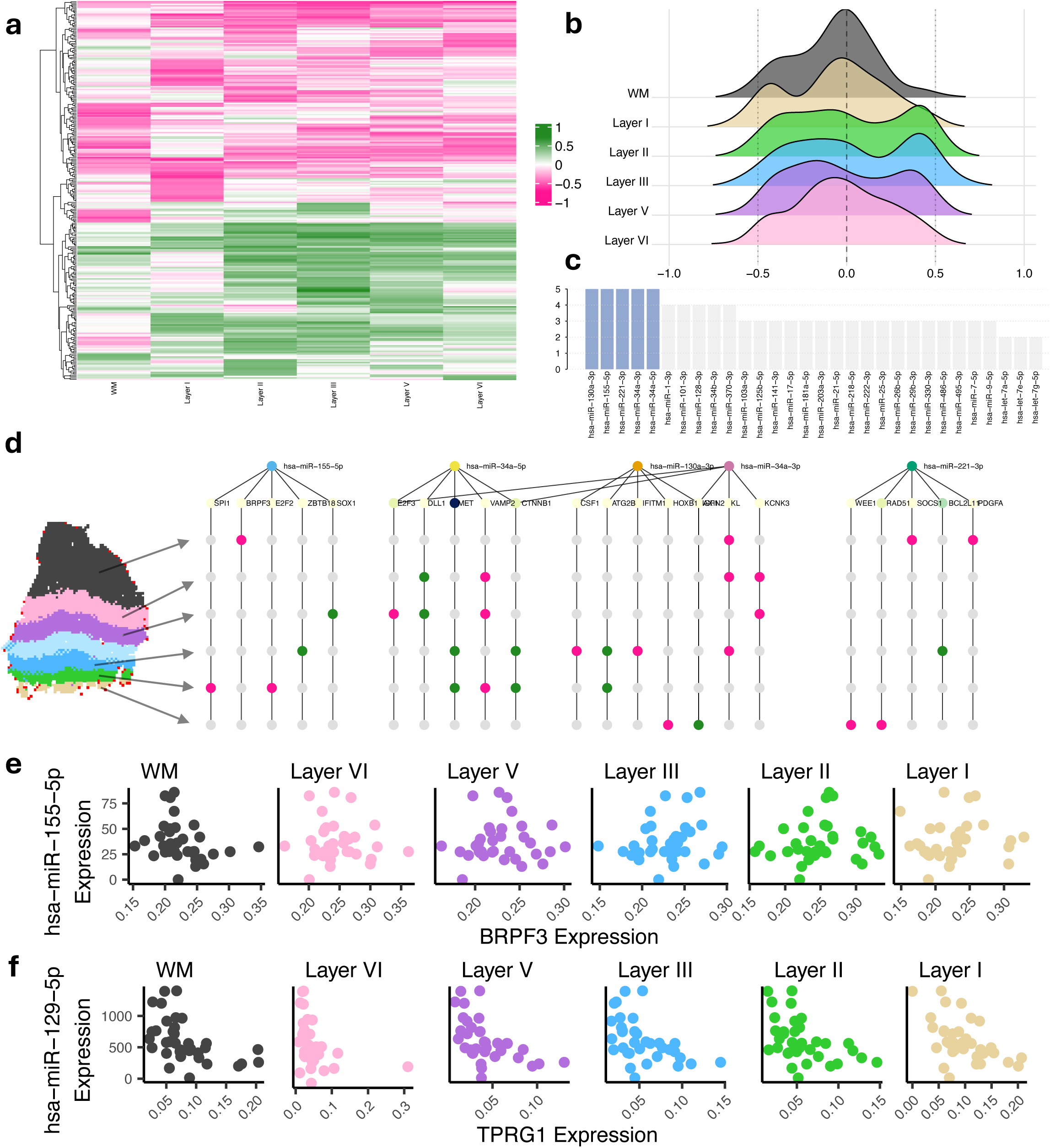
Layer-specific miRNA regulation **a,** Heatmap displaying Spearman correlation coefficients for miRNA-mRNA interactions across all spatial domains. **b,** Ridge plot illustrating the distribution of correlation coefficients throughout spatial domains. **c,** Quantification of high-correlation interactions per miRNA. **d,** Network diagram showing five miRNAs (colored circles at top: hsa-miR-155-5p, hsa-miR-34a-5p, hsa-miR-130a-3p, hsa-miR-34a-3p, and hsa-miR-221-3p) connected to their target genes (labeled below each miRNA). Each vertical line represents a miRNA-mRNA interaction, with multiple nodes along each line representing different spatial domains. Nodes are colored green for strong positive correlations (>0.4), pink for strong negative correlations (<-0.4), and grey where no strong correlation exists between the miRNA and mRNA in that particular spatial domain. **e,** Scatter plots correlating miR-155 expression with BRPF3 expression across spatial domains. **f,** Scatter plots correlating miR-129 expression with TPRG1 expression across spatial domains.

Comparison of correlation distributions across spatial domains revealed distinct shifts, strongly suggesting distinct layer-specific patterns of miRNA regulation (**Fig. 5b**). To identify miRNAs with substantial regulatory impact, we quantified high-correlation miRNA-mRNA interactions (|R|>0.4) for each miRNA across all domains. We identified five miRNAs that exhibited five or more high-correlation interactions with mRNAs across different spatial domains (**Fig. 5c**). Notably, this group included miR-155 and miR-34a, which have previously been linked to neuroinflammatory processes and neurodegeneration, respectively^65,67,68^. All five miRNAs demonstrated substantial layer-specific distribution patterns, often showing strong regulatory activity (either positive or negative) in only one cortical domain (**Fig. 5d**).

Our analysis revealed a complex regulatory landscape with two distinct patterns of miRNA activity across cortical layers. Some miRNAs function as global regulators with consistent activity throughout all cortical layers, while others display pronounced layer-specific regulation. This spatial specificity is exemplified by miR-155, which predominantly regulates BRPF3 expression in white matter (**Fig. 5e**), contrasting with miR-129, which maintains strong regulatory control of TPRG1 expression throughout all cortical domains (**Fig. 5f**).

In sum, our findings suggest that miRNA-mediated post-transcriptional regulation contributes significantly to maintaining the unique molecular signatures of different cortical layers, adding another dimension to our understanding of the mechanisms governing cortical lamination.

## DISCUSSION

Our findings reveal extensive layer-and region-specific transcriptional dysregulation at both spot and single-cell resolution, highlighting a fundamental limitation of conventional single-cell RNA-sequencing approaches that fail to capture spatial context. By preserving tissue position information, we demonstrate that disease-associated gene expression patterns are highly dependent on anatomical location, with transcriptional dysregulation signatures varying dramatically across cortical layers and between brain regions.

Specifically, we identified a distinct stress response signature upregulated in Parkinson’s disease compared to dementia with Lewy bodies, which is most prominent in Layer V neurons. Analysis of the transcriptional microenvironment surrounding stressed cells unveiled remarkably enhanced metabolic and transcriptional activity as well as upregulated apoptotic and protein degradation pathways at the stress epicenter and its immediate neighboring cells, while developmental and regenerative pathways were progressively restored at increasing distances from the stress epicenter. Additionally, known PD risk genes were significantly upregulated in the stress epicenter, with expression gradually diminishing in more distant cells. Further mechanistic studies examining this spatial stress response gradient will be essential for understanding the selective vulnerability of these neurons and may provide novel therapeutic targets for intervention.

In COVID-19 patients, our spatial analysis revealed significant peripheral immune cell infiltration into the brain parenchyma, particularly seen in the white matter of the anterior cingulate cortex. This infiltration was characterized by distinct transcriptional signatures in microglia, endothelial cells, and astrocytes, suggesting compromised blood-brain barrier function and coordinated neuroinflammatory responses. Neuropathological studies have described these focal immune cell infiltrations not only in the subcortical white matter, but also in the brain stem^51^. These findings align with and extend prior research documenting blood-brain barrier dysfunction and persistent systemic inflammation in long-COVID patients^8^. Similar to our findings in PD, analysis of immune hotspots revealed transcriptional gradients with inflammatory pathways dominating near these hotspots while neuroprotective mechanisms were suppressed. These spatial patterns may help explain the neurological manifestations of COVID-19 and warrant further investigation to determine their persistence in long-COVID patients and potential contribution to lasting neurological symptoms. Future comparative studies with other neurotropic viruses will be crucial to distinguish SARS-CoV-2-specific neuropathology from general viral neuroinflammatory mechanisms and to elucidate the precise pathways through which these immune infiltrates may damage neural tissue.

In total, we present one of the first single-cell spatial transcriptomics atlases of the human brain to date, encompassing 40 tissue samples across two distinct cortical regions from patients with various neurodegenerative conditions and severe COVID-19. As one of the first and most comprehensive human brain single-cell spatial datasets to date, this atlas provides a valuable resource for the scientific community, both to validate existing hypotheses and discover novel biological insights, as well as for the development of scalable analysis methods for spatial transcriptomics data.

While spatial transcriptomics at single-cell resolution provides unprecedented insights into tissue biology in health and disease, several limitations and challenges remain associated with this technology. Similar to single-cell RNA-seq, established differential expression analysis is influenced by pseudo-replication bias, where inter-replicate variation is not adequately accounted for^69^. This issue is further magnified in single-cell spatial transcriptomics due to the increased number of cells per donor. To address this concern, we provided information at the donor level for our major results, demonstrating that expression signatures are consistently dysregulated across multiple donors. Furthermore, there are currently no comprehensive studies investigating the influence of post-mortem interval on spatial transcriptomics data quality. Consequently, we cannot exclude the possibility that post-mortem changes may affect the biological signals observed. Additionally, large-scale multimodal studies will be necessary to validate and extend our spatial findings, particularly to investigate potential differences related to sex and age.

While gene expression can be measured at the single-cell level and even with spatial resolution, single-cell miRNA quantification remains challenging, as most techniques rely on poly-A tails, which miRNAs lack. Although single-cell approaches exist^70^, they currently offer limited sensitivity. To address this, we measured miRNAs at the bulk level and aggregated spatial single-cell data at domain-and patient-level pseudobulk resolution, enabling direct comparisons. By prioritizing high-confidence, experimentally validated targets over weak or predicted ones, we refined our focus while acknowledging a degree of selection bias. Notably, the miRNAs we identified align with known PD-associated candidates, reinforcing their biological relevance. While some are also linked to other diseases, such as cancer^71,72^, their layer-specific regulation in the brain points to a distinct functional role in PD pathology.

## METHODS

### Human tissue processing and Stereo-seq Spatial Transcriptomics

Kryopreserved tissues were selected retrospectively from the biobank of the Institute of Neuropathology of Saarland University based on clinical parameters. Eligible tissues were subsequently cut to approx. 0.9*0.9cm sized blocks and embedded in Tissue Tec (Leica, Wetzlar, Germany). RNA integrity was checked by isolation of RNA from 5-10 10µm sections using the miRNeasy Tissue/Cells Advanced Mini Kit (Qiagen, Hilden, Germany) and RNA 6000 Nano Bioanalyzer Kit (Agilent, Santa Clara, CA, USA). Only tissues with RIN>7 were used for spatial transcriptomics using the Stereo-seq Transcriptomics T Kit (BGI, Shenzhen, China) according to the manufacturers protocol. In short, a 10µm section of the tissue was placed on the Stereo Seq chip and fixated in ice-cold methanol for 30 min. Subsequently, nuclei were stained using the fluorescent DNA stain of the Qubit ssDNA kit (Thermo Fisher Scientific, Waltham, MA, USA). The chips were scanned in a MOTIC microscope (MOTIC, Hongkong, China) with 10x resolution in FITC channel to generate an image containing the spatial information of the cells on the grid background of the chip enabling read annotation in a spatial context at the end of the experiment. Afterwards, tissue was permeabilized for 18 min to release RNA from the cells and enabling the binding to the spatially barcoded DNBs on the chip. Next, mRNAs bound to the chip were reverse transcribed into cDNA for 3 hours at 42°C and then released from the chip at 55°C over night. The following day, cDNA was collected, PCR amplified and purified using SPRI select magnetic beads (Beckman Coulter, Brea, CA, USA). CDNA quality was checked using Bioanalyzer High Sensitivity chip (Agilent, Santa Clara, CA, USA). For each sample, 20ng cDNA were used for library construction including a cDNA fragementation step, a PCR amplification step with introduction of the sample barocode, and subsequent purification of the final library product using SPRI select magnetic beads (Beckman Coulter, Brea, CA, USA). Library size was checked using Bioanalyzer High Sensitivity chip (Agilent, Santa Clara, CA, USA). Libraries were sequenced on a DNBSEQ-T10 sequencer by MGI Tech. Riga using a paired end 75 bp strategy.

### Immunohistochemistry

Human brain tissue (medial frontal cortex and anterior cingulate cortex, each with cortex-associated white matter) adjacent to tissue processed for single-cell spatial transcriptomics was subjected to immunohistochemistry (IHC). Seven µm sections of frozen tissue were cut and fixed in 4% formalin for 1 hour. Peroxidases were blocked by incubation in 1% H_2_O_2_ for 15 minutes at room temperature. Heat antigen retrieval was performed by steaming at 98°C in target retrieval solution pH 6.1 (Dako, Carpinteira, CA, USA #S1699) for 30 minutes. Sections were allowed to cool down at room temperature. Following antigen retrieval, sections were incubated for 45 minutes at room temperature with the mouse monoclonal anti-CD163 antibody clone 10D6 (Leica, NCL-L-CD163; 1:200). The antibody was diluted in Dako REAL antibody diluent #S2022. After three washes with wash buffer (Dako #S3006), the Dako REAL EnVision HRP kit (#K5007) was used for the visualization of the antibody reaction according to the manufacturer’s instructions. Sections were counterstained with Mayer’s hemalum (Sigma-Aldrich #1.09249). After dehydration, coverslips were mounted with Entellan (Merck #1.07961). Images were acquired with an Olympus BX 40 microscope, equipped with an Olympus SC30 digital microscope camera using the Olympus cellSens software.

### Bioinformatics and quality control

Prior to computational analysis, we conducted comprehensive quality assessment of each tissue section. This included visual inspection of overall tissue morphology and integrity, as well as evaluation of spatial expression patterns for established anatomical domain markers MBP, SNAP25 and PCP4. Samples exhibiting significant tissue degradation or displaying aberrant localization of these canonical markers relative to their expected anatomical distributions were excluded from downstream analyses to ensure data quality and biological relevance.

Outputs from the Stereo-seq Analysis Workflow (SAW) pipeline were converted into Seurat v5.2 objects for both bin200 (representing 200×200 Stereo-seq spots) and cellbin formats. Conversion was performed using the io.stereo_to_anndata function from the Stereopy package (v1.1.0), followed by implementation of the h5ad2rds.R script provided by BGI to generate Seurat objects.

For cellular analyses, cell bins were defined through image-based cell segmentation of the corresponding single-stranded DNA (ssDNA)-stained images. Quality control was applied to the cellbin samples using the following criteria: a minimum of 200 expressed genes per cell, a minimum of 300 unique molecular identifier (UMI) counts per cell, and a maximum mitochondrial DNA content threshold of 15%. Cells failing to meet these quality metrics were excluded from downstream analyses.

### Spatial Domain Clustering and Annotation

Bin200 Seurat objects were converted to SingleCellExperiment objects and preprocessed using the *spatialPreprocess* function from the BayesSpace library with 2,000 highly variable genes (HVGs). Cross-sample integration across both anatomical regions was achieved using the *RunHarmony* function from the Harmony package^73^ with sample identifiers as grouping variables. Spatial coordinates were adjusted to prevent overlapping between different spatial samples. BayesSpace clustering was then performed on the integrated dataset using the *spatialCluster* function with the following parameters: 10 clusters (q=10), 40 dimensions (d=40), spatial smoothing parameter of 3 (gamma=3), 10,000 MCMC iterations (nrep=10000), and 100 burn-in iterations (burn.in=100). The resulting spatial domains were annotated based on established canonical markers from previous studies^34,35,41^

### Cell Type Annotation by Reference-Based Label Transfer

Cell type annotation was performed using a reference-based label transfer approach. All cellbin samples were converted to the Scanpy framework^74^. The Seattle Alzheimer’s Disease (SeattleAD)^43^ dataset served as the reference for label transfer, which was implemented using the destVI (DEconvolution of Spatial Transcriptomics data using Variational Inference) method^42^.

The annotation process was conducted in two sequential rounds. In the first round, broad cell type categories were assigned, including various glial cell populations and neurons categorized as either excitatory or inhibitory. Subsequently, a second round of annotation was performed specifically on neuronal populations to achieve fine-grained classification of excitatory and inhibitory neurons into their respective subtypes.

For each cell, the cell type assignment was based on the highest probability score determined by the destVI algorithm. All annotation procedures followed the established destVI tutorial guidelines. To improve gene selection in noisy spatial transcriptomics data, we added a specified set of canonical cell type markers to the scvi gene set. We used the following parameters:

*Reference subsample proportion*: 0.4; *n_hvg_sc*: 3000; *n_hvg_st*: 20000; *n_epochs_sc*: 300; *n_epochs_st*: 10000; *batch_size*: 512

Model training was performed on one NVIDIA H100 GPU.

### Neighborhood Analysis and Spatial Gene Gradients

Signature scores were calculated for all cells using the *AddMetaData* function in Seurat. Cells with signature scores in the top n% were designated as signature hotspots. To characterize the spatial organization relative to these hotspots, we computed the Euclidean distance from each cell to its nearest signature hotspot using the spatial coordinates within each sample. The resulting distance matrices were subsequently concatenated across all samples.

To define spatial neighborhoods, cells were binned according to their distance from signature hotspots, with Neighborhood1 comprising hotspot cells and cells within 50 μm of a hotspot. Subsequent neighborhoods (Neighborhood2-8) were defined at 50 μm increments (i.e., Neighborhood2: 50-100 μm, Neighborhood3: 100-150 μm, etc.). For downstream analyses, we focused on these eight neighborhoods (Neighborhood1-8, covering distances up to 400 μm from hotspots).

To identify genes with expression patterns that vary systematically across these spatial neighborhoods, we applied the scITDG package (https://github.com/YandongZheng/scITDG). This analysis enabled the identification of spatially dysregulated genes across neighborhoods. The dysregulated genes were further clustered into k=6 modules based on their expression patterns, and trajectory plots were generated to visualize the spatial gradients of gene expression.

### Differential Expression Analysis

Differential expression analysis throughout the manuscript was performed using the Model-based Analysis of Single-cell Transcriptomics (MAST)^75^ with latent variables *nCount_RNA* and *PatientID*

### Pathway Analysis

We performed all pathway analyses using the Enrichr^76^ library with gene sets *GO Biological Process* 2023 and *KEGG 2021 Human*.

### miRNA-mRNA linkage analysis

For the analysis, we employed miRMaster 2.0^77^ using its default parameters on the provided datasets. This tool performed sequence alignment against the human genome and mapped reads to known miRNAs using miRbase^78^. The alignment and mapping steps were conducted with Bowtie^79^, applying the settings “-m 100 —best —strata” to generate raw miRNA count data. Additionally, we collected the alignment and mapping details generated during processing. For downstream analysis, we allowed a misclassification rate of 1.

To ensure comparability across samples, raw counts were subjected to rpmm-normalization. We then applied filtering at both the sample and feature levels. Specifically, only samples with over one million reads aligned to the human genome were retained. Features were filtered based on the criterion that at least one disease group within at least 10% of the samples exhibited a raw count of five or higher. Following this filtering step, all 35 samples (matching the spatial transcriptomics samples passing initial QC) remained, along with 1277 of 2656 miRNAs were retained for further analysis. For each spatial domain, we computed Spearman rank correlation coefficients between miRNAs and mRNAs (pseudobulk expression per spatial domain and patient using *AggregateExpression* in Seurat) with known functional relationships, as defined in miRTarBase^66^. The resulting matrix was subsequently filtered to contain only miRNA-mRNA interactions having strong correlation (|R| > 0.4) in one spatial domain and show variable correlations across domains (SD > 0.15).

### Statistics and reproducibility

We used the following package versions for our analysis:

R v4.4.2, python v3.10.14, scvi-tools v1.2.0, pytorch v2.4.1, scanpy v1.10.3, numpy v1.26.4, pandas v2.2.3, Steropy v1.1.0, Seurat v5.2.1, SeuratObject v5.0.2, enrichR v3.4, harmony v1.2.3, igraph v2.1.4, MAST v1.32.0, tidyverse v2.0.0, ggplot2 v3.5.1, ggraph v2.2.1, ggridges v0.5.6, pheatmap v1.0.12, dplyr v1.1.4, tibble v3.2.1, SingleCellExperiment v1.28.1, BayesSpace v1.16.0, ggrepel v0.9.6 Raw p-values were adjusted for multiple testing bias using Benjamini-Hochberg FDR correction method.

## Data availability

The sequencing dataset generated in this study will be made available in the NCBI Sequence Read Archive (SRA) at the day of publication.

## Code availability

The code used to generate our results will be made available on Github at the day of publication.

---

**Supplementary Fig. 1.**
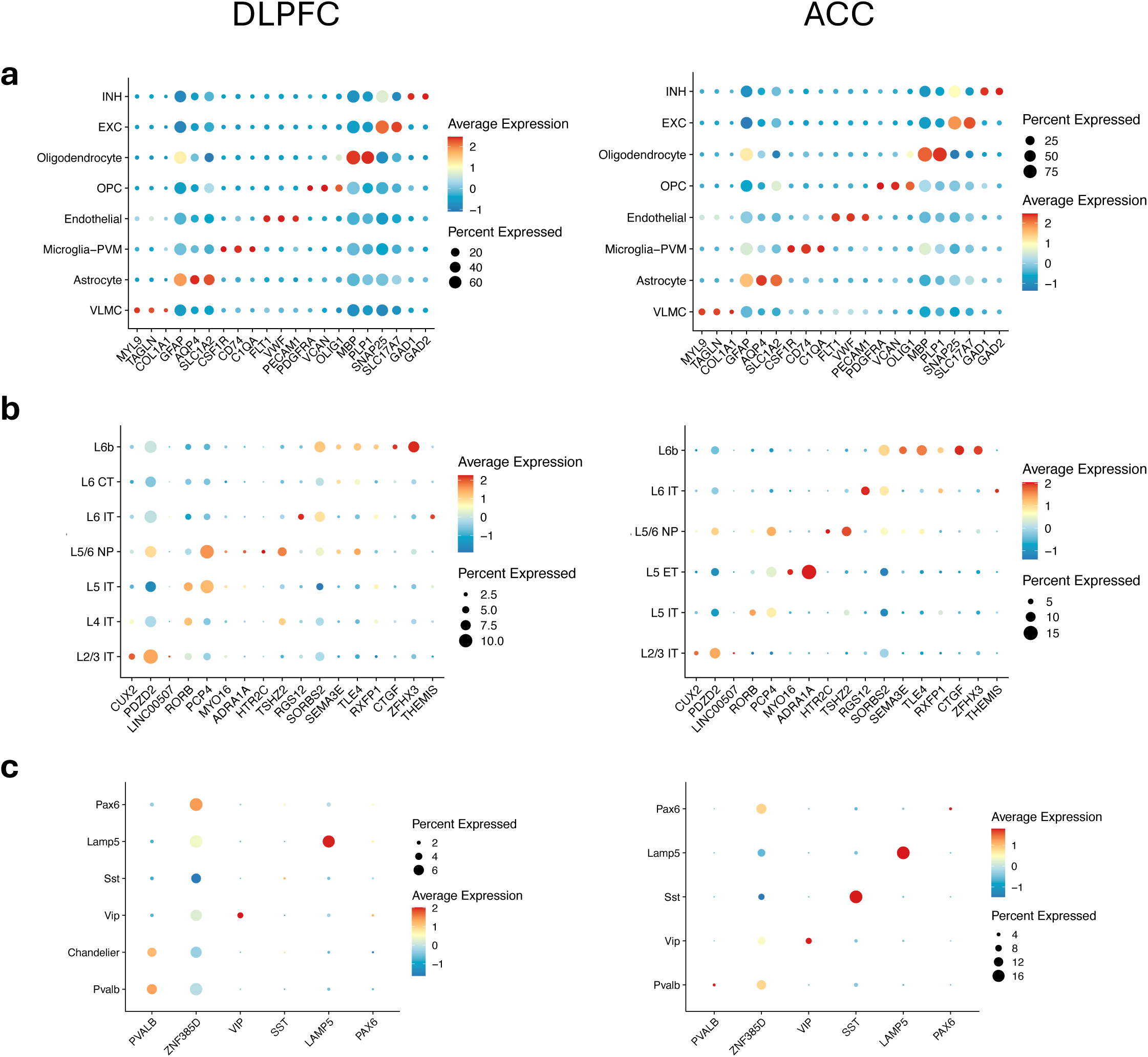
a-c, Dotplots showing relative expression of marker genes for (a) broad celltypes, (b) excitatory neuron subtypes and (c) inhibitory neuron subtypes

**Supplementary Fig. 2.**
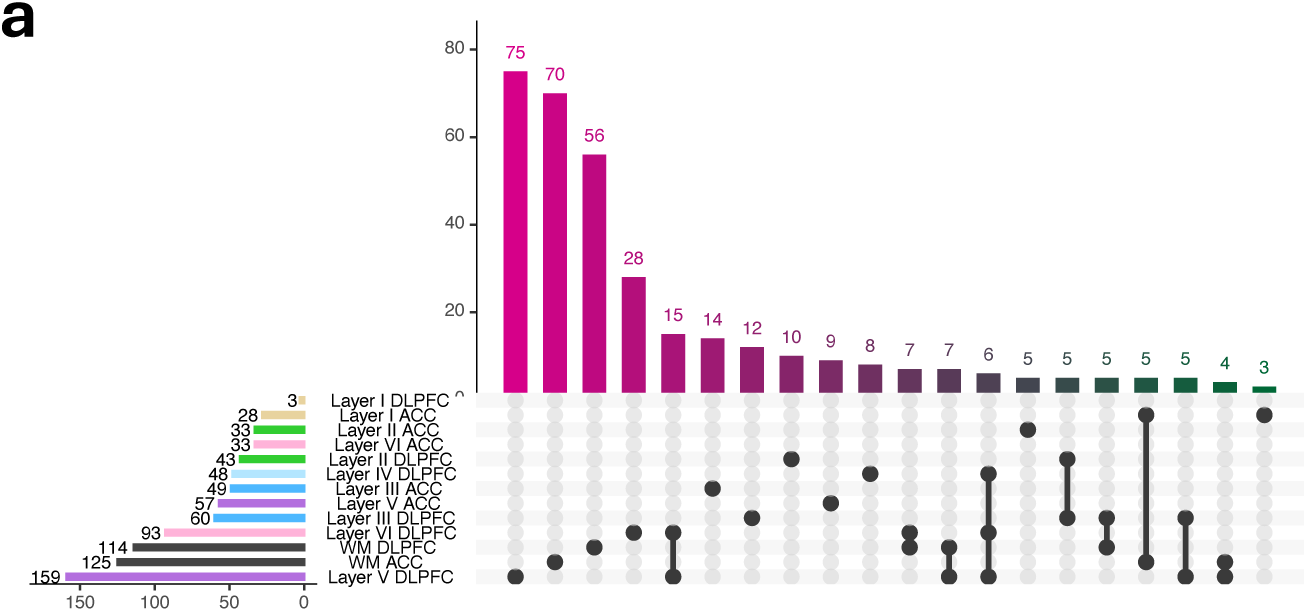
a, Upset plot visualizing the intersection of COVID-specific differentially expressed gene (DEG) sets across spatial domains and cortical regions.

**Supplementary Fig. 3.**
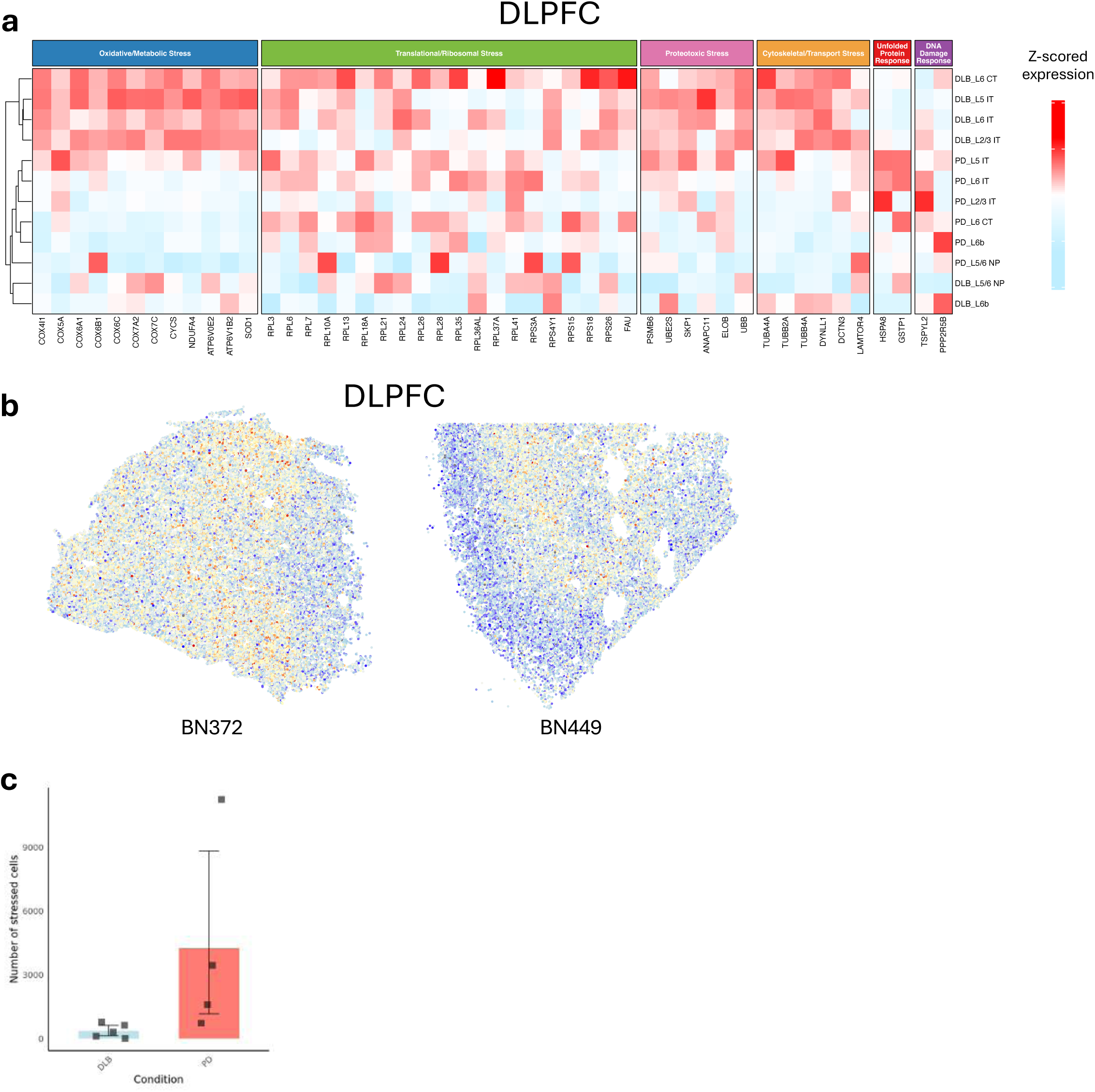
a, Heatmap displaying z-scored average expression profiles in DLPFC excitatory neurons, comparing PD and DLB patients. **b,** Spatial visualization contrasting stress signature scores between representative PD and DLB samples from the DLPFC. **c,** Boxplot showing the number of stressed cells per patient compared between PD and DLB

**Supplementary Fig. 4.**
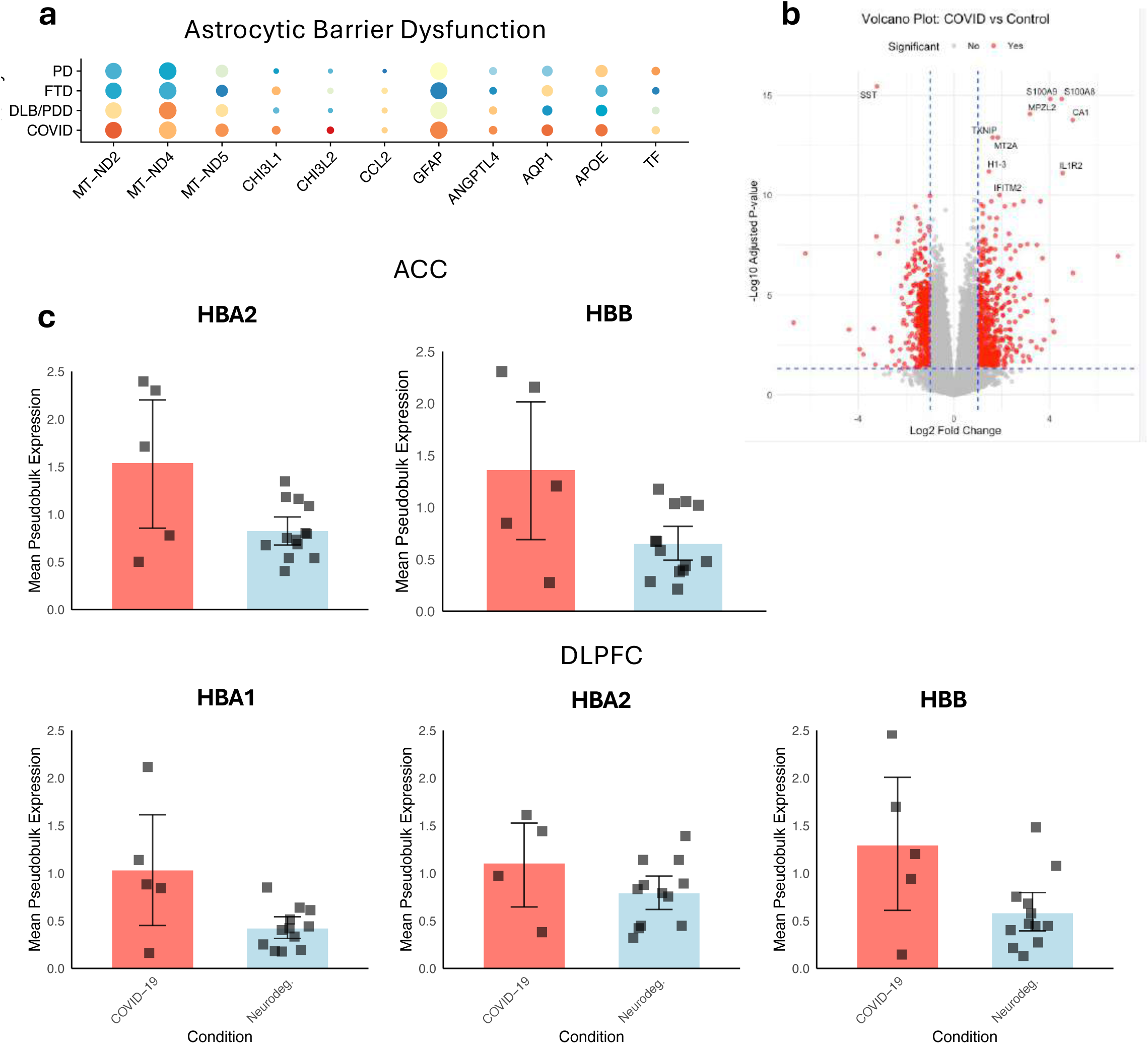
a, Differential gene expression in astrocytes across all four diseases, specifically within the ACC white matter **b,** Volcano plot showing significantly dysregulated genes in a reference bulk RNA-seq dataset **c,** Boxplot comparing Hemoglobin (HBA1, HBA2 and HBB) gene pseudobulk expression per sample in the ACC and DLPFC white matter between COVID-19 and the pooled cohort of all three neurodegenerative conditions.

**Supplementary Fig. 5.**
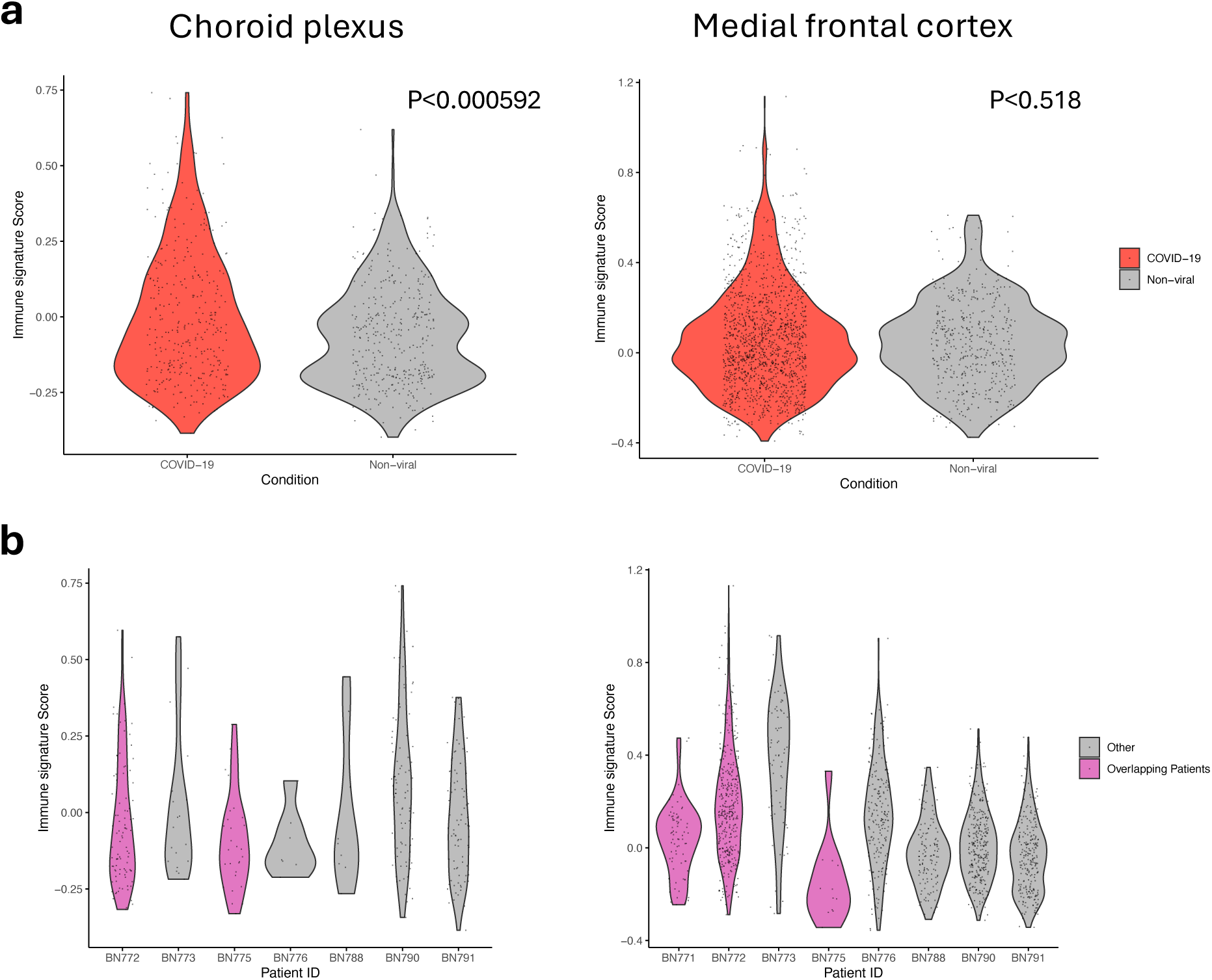
a, Violin Plot showing immune signature scores in Macrophages/Microglia of the choroid plexus and the medial frontal cortex in a COVID-19 single-cell dataset comparing between COVID-19 and non-viral controls **b,** Violin Plot showing immune signature scores in Macrophages/Microglia of the choroid plexus and the medial frontal cortex in a COVID-19 single-cell dataset comparing between COVID-19 patients. Pink violins correspond to patients overlapping with our current study.

**Supplementary Fig. 6.**
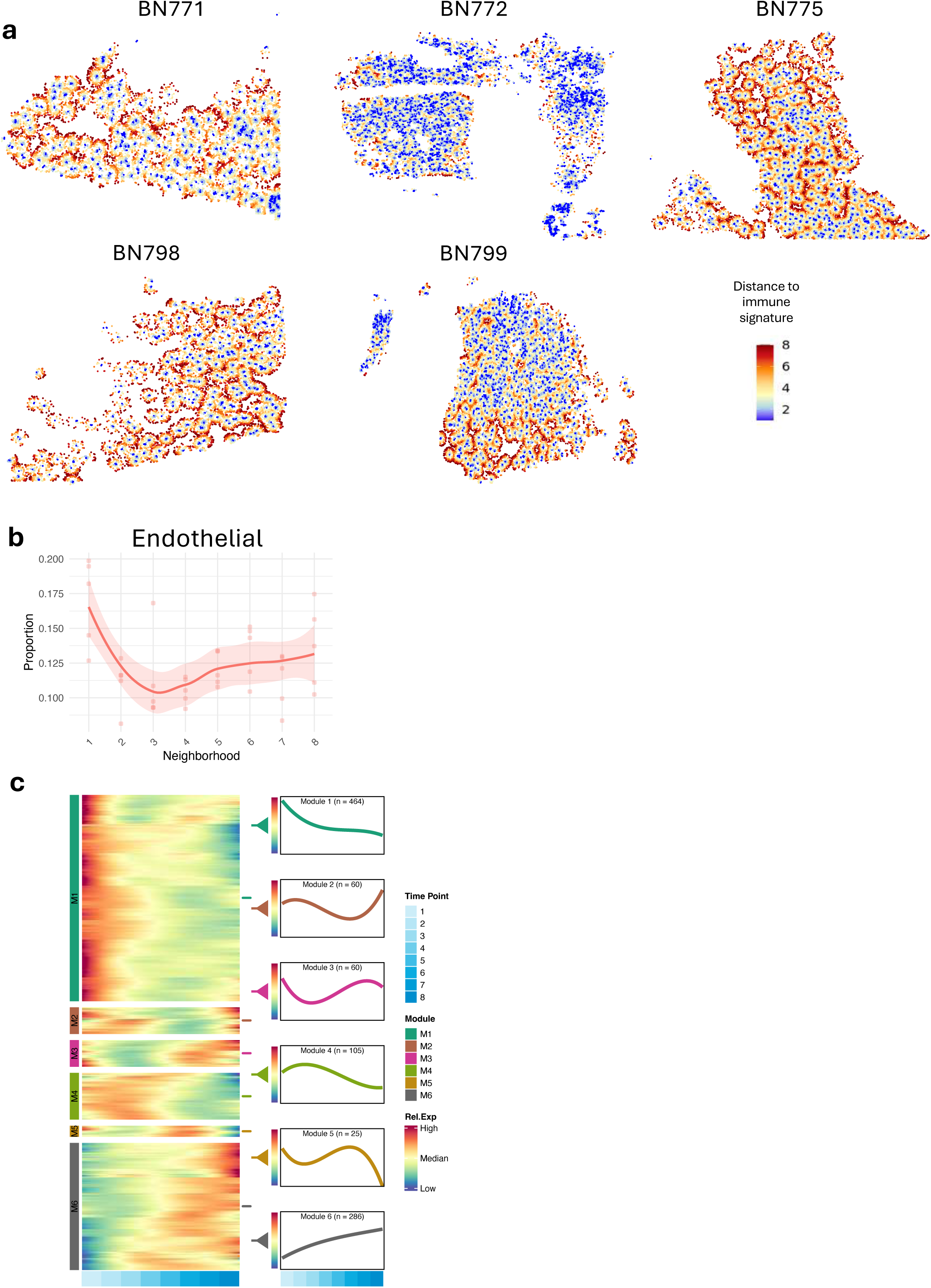
a, Spatial visualization highlighting cells in neighborhoods 1-8, with all other cells omitted for clarity, for all PD samples not shown in the main figure. **b,** Cell type proportions of endothelial cells across neighborhoods 1-8 within immune hotspots. **c,** Heatmap illustrating the expression patterns of modules 1-6 across neighborhoods 1-8.

